# Coupling fibroblast mechanotransduction signaling to tissue growth in a multiscale model of skin expansion

**DOI:** 10.64898/2026.07.31.742141

**Authors:** Laura Nunez-Alvarez, Joanna K. Ledwon, Parul Rai, Kathryn Reisner, Tianhong Han, Luis Solorio, Arun Gosain, Adrian Buganza Tepole

## Abstract

Skin growth and remodeling underlies health, disease, and treatments such as tissue expansion (TE). The mechanotransduction pathways in dermal fibroblasts are increasingly well characterized, and tissue-level growth has been described phenomenologically, but coupling between cell-level signaling and tissue-level growth remains poorly understood. We develop a dermal fibroblast signaling network through extensive literature data curation, comprising 151 reactions among 96 nodes. The inputs are mechanical stretch and eight ligands (TGF*β*, PDGF, FGF, IL1, IL6, TNF*α*, AngII, ET1); outputs of interest span ECM-enzymes (proMMP1/2/9, MMP1/2/9), ECM proteins (CImRNA, collagen I, fibronectin), and fibroblast activity (*α*SMA, proliferation). Implemented as a logic-based ODE system, the network reproduces 82% of the calibration dataset and agrees with independent validation data. Sensitivity analysis reveals a tension-dependent regulation of signaling: at baseline tension, outputs are governed by many boosters (nodes that positively influence downstream targets) and one dominant brake, LATS1/2, whereas at high tension control consolidates and new, tension-specific regulators such as integrin (ITGB1) emerge. Multiple pathway axes converge on a few central regulators, producing pronounced crosstalk, most notably between TGF*β* and mechanical tension. Finally, linking the collagen outputs to a tissue-level growth formulation yields a bidirectional mechanical–biochemical coupling that reproduces tension-induced skin growth measured in a porcine TE model. This framework establishes a comprehensively calibrated dermal fibroblast signaling network coupled to tissue-level growth, opening opportunities for targeted TE interventions.

## Introduction

Skin growth and remodeling are processes regulated by mechanical stimuli and biochemical signaling. A clinically relevant example is the reconstruction of large skin defects using tissue expansion (TE), in which an expander is inserted beneath the skin and gradually inflated to promote growth in response to a sustained tensile stimulus [1]. External stretching is well established to activate mechanoresponsive pathways, leading to the production of extracellular matrix (ECM) [2, 3] and cell proliferation [4, 5], both of which are key markers of dermal tissue growth. This is possible because mechanosensing elements, such as integrins, ion channel receptors, and growth factor receptors, work together with signaling molecules to eventually trigger gene expression in the process known as mechanotransduction [6]. Mechanotransduction allows dermal fibroblasts to maintain tissue homeostasis under normal conditions and regulate growth under external mechanical stimulus [7]. If abnormal conditions persist, they may transition to a profibrotic state, leading to excessive collagen production [8]. In both cases, fibroblast-mediated ECM remodeling, downstream of mechanical cues, alters tissue composition and, consequently, its mechanical stress state, creating a feedback loop between the mechanical environment and the biochemical response.

Because mechanical stimuli and biochemical signaling are tightly coupled, a complete description of skin growth and remodeling (G&R) under TE requires understanding not only the tissue-level mechanical response but also the feedback loop connecting the mechanical environment to the underlying cellular signaling cascade. Despite significant progress in characterizing individual mechanotransduction elements, their coupled intracellular dynamics remain poorly characterized. Moreover, how this cell-level signaling activity feeds back to drive tissue-level growth remains unclear. Existing calibrated models of TE have described growth with a phenomenological, tissue-level perspective [9, 10, 11]. They have proven useful for predicting outcomes within that scope. However, because they do not represent the biochemical response explicitly, they are fundamentally limited to tissue-level predictions. For example, they cannot model interventions that act at the cellular level such as pharmacological treatments. A model that explicitly couples the mechanical environment to the biochemical signaling cascade would potentially enable the design of biologically targeted interventions and open the door to patient-specific calibration using both tissue-level as well as cell-level data.

Multiple mechanosensitive pathways contribute to skin remodeling, and their interactions are highly context-dependent. Under baseline tension, canonical mechanosensors such as PIEZO1 and integrins are expressed at relatively low levels in dermal fibroblasts [12, 13, 5]. PIEZO1 mediates calcium influx [12], whereas integrins promote the formation of focal adhesion kinase (FAK) complexes, which coordinate cell survival, proliferation, and motility through FAK-Src–dependent signaling [14]. When external tension increases beyond homeostasis, PIEZO1-mediated calcium influx promotes fibroblast proliferation, migration, and collagen synthesis [8]. Integrin-FAK activation under high tension stimulates downstream RhoA/ROCK signaling [15] and interfaces with important effectors of the Hippo pathway [8], thereby regulating myofibroblast activation and ECM production. Additional mechanotransductive axes, including the MAPK/ERK pathway, further enhance collagen synthesis and support ECM homeostasis [8] in part by potentiating TGF*β* signaling and increasing the expression of structural proteins such as collagen I (CI), collagen III (CIII), fibronectin (FN), and connective tissue growth factor (CTGF) [16].

Importantly, these signaling axes do not function independently. PI3K acts as a central integrator of diverse biochemical and mechanical cues: it is activated downstream of TGF*β*, PDGF, and FGF, as well as inflammatory mediators including TNF*α* and adhesion-dependent signaling through integrins [17, 18, 19, 15]. Such convergence highlights how fibroblasts coordinate multiple upstream inputs to regulate cytokine expression, survival, and matrix remodeling in mechanically dynamic environments.

A second layer of complexity comes from the fact that individual regulators often carry diverse and sometimes opposing, biochemical functions independent of mechanical context. The matrix metalloproteinase (MMP) family exemplifies this: several MMPs (MMP1, MMP2, MMP14) are canonical interstitial collagenases traditionally viewed as collagen-degrading enzymes [20], yet a subset (MMP2 and MMP9) also cleave latent TGF*β* stored at the fibroblast surface, thereby promoting the collagen synthesis [21]. This functional heterogeneity, where the same protein contributes to both matrix turnover and its promotion is difficult to disentangle from pathway-by-pathway analysis and, together with the tension-context effects noted above, motivates a network-level framework.

To model cellular signaling activity, several approaches can be used. Pure mechanistic models, based on biochemical kinetics or the mass-action law, offer detailed descriptions of molecular interactions but require calibration and measurement of many parameters [22]. This poses a data collection challenge, particularly when dealing with complex networks involving multiple pathways and species, for which quantitative data may be incomplete or unavailable. On the other end of the spectrum, simpler Boolean models provide qualitative insight into network logic but often lack continuity in both state (i.e., on/off of multi-state discrete systems) and time, as observed in biological systems [23]. Logic-based dynamic (ODE) models offer a compromise between these extremes. They incorporate signaling network topology and logic of Boolean models while introducing continuous-valued dynamics that better reflect biological signaling. This intermediate formulation enables the simulation of time-dependent responses without requiring detailed kinetic parameters, making it a practical and biologically relevant modeling approach. Logic-based ODE models have been widely applied to study ECM remodeling [24, 25, 26], and have been coupled with tissue-level growth dynamics in cardiac applications [27, 28], which is highly relevant to our TE scenario.

In this work, we develop a multiscale model of fibroblast-driven skin G&R under TE. At the cell level, we adopt a logic-based ODE framework to describe the coupled dynamics of key mechanotransduction pathways in response to mechanical loading. The fibroblast signaling network is informed by a comprehensive literature review and data curation. At the tissue level, the predicted biochemical activity is coupled to a growth formulation based on continuum mechanics [29]. The multi-scale model is validated with experimental data on skin growth during TE from a porcine animal model [9].

## Materials and Methods

### Network structure

We constructed a dermal fibroblast signaling network by adapting a previously published cardiac fibroblast network [24]. We removed nodes and interactions specific to the cardiovascular system, retained nodes with evidence in dermal fibroblasts, and added three signaling axes relevant to skin G&R. The resulting network connects nine inputs to eleven outputs through 96 nodes and 151 reactions. Inputs consist of one mechanical stimulus (stretch) and eight ligands: IL1, IL6, TGF*β*, PDGF, ET1, AngII, TNF*α*, and FGF [30, 31, 32, 33, 34, 35, 36, 37, 38]. For TGF*β* in particular, we defined a dedicated *TGFβ infusion* input node to represent upregulated TGF*β* signaling independent of mechanical stretch, allowing direct perturbation of the pathway separately from mechanically induced activation of latent TGF*β* stored in the ECM.

The eleven outputs of interest represent downstream markers of fibroblast-driven G&R: two cell-level processes (*α*SMA expression, and proliferation), two ECM proteins (collagen type I, and fibronectin), and seven intermediate regulators of ECM (proMMP1/2/9, MMP1/2/9, and CImRNA). Collagen type III (CIII) and its transcript (CIIImRNA) share identical upstream regulatory logic with CI and CImRNA, respectively, and are therefore not included as separate outputs to avoid redundancy. A schematic of the network is shown in Fig. 1.

**Figure 1:**
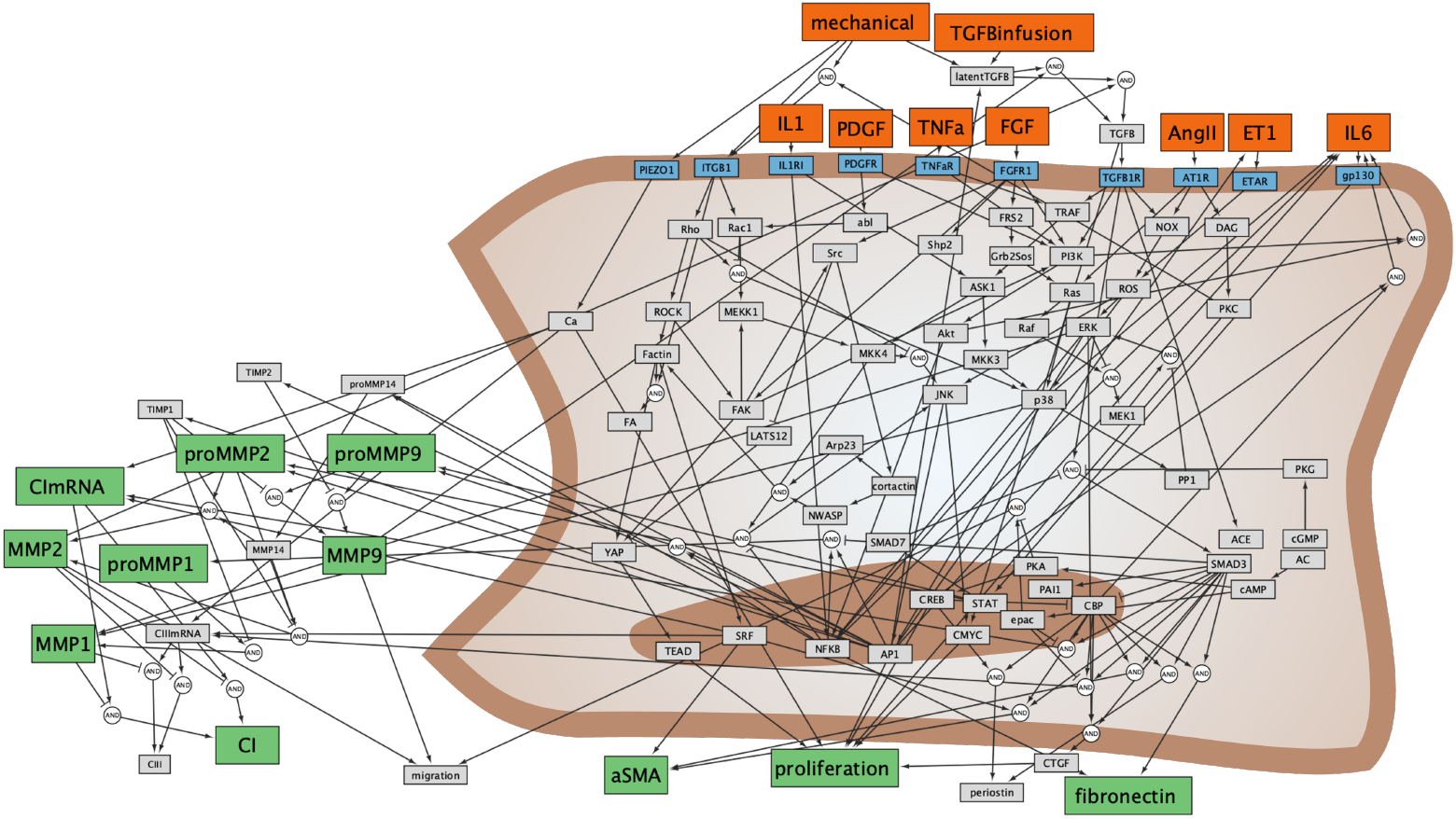
Signaling network describing dermal fibroblast responses to biochemical and mechanical stimuli. External inputs (orange) represent cytokines and mechanical cues that activate membrane receptors (blue). Intracellular nodes correspond to intermediate signaling molecules (grey). Transcription factors localized to the nucleus integrate upstream signals and regulate downstream phenotypic responses. Model outputs (green) represent key fibroblast phenotypes, including collagen I/III production, matrix metalloproteinase activity, proliferation, and *α*SMA expression. Nodes grouped on the left side form a collagen remodeling module that captures the balance between collagen synthesis and degradation. Edges indicate activating or inhibitory regulatory interactions compiled from literature.

We then conducted an extensive literature review to refine the reactions linking all the nodes of the network. We required each interaction to be supported by at least two experimental studies, resulting in 151 reactions in total. Of these, 85 were validated in dermal fibroblasts (78 in human-derived lineages and 7 in mouse models), 31 in fibroblast-like cells (29 cardiac, 2 lung), and 34 in mammalian fibroblasts more broadly. For pathway crosstalk interactions, we included reactions that were well characterized in other cell types but supported by evidence in fibroblast-related cells. Cell-free studies were used to support extracellular reactions [24]. The logical network was then translated into an ODE system using Netflux software [39], as detailed in Suppl. 2.

Mechanical stretch is sensed through two complementary entry points. Integrin clustering under load assembles focal adhesion complexes, activating FAK and initiating downstream RhoA/ROCK signaling [40, 41]. In parallel, stretch increases the open probability of PIEZO1, driving calcium influx [12].

Downstream of these entry points, mechanotransduction intersects with growth factor and inflammatory signaling through shared nodes. Growth factor pathways (FGF, PDGF) activate Ras-MAPK cascades, whose amplitude and duration are modulated by focal adhesion signaling and cytoskeletal tension [38, 34]. Inputs AngII and ET1 introduce redox-dependent regulation through ROS [36, 35], while inflammatory cues converge via ASK1-dependent activation of p38 and JNK [8]. The network architecture thus allows mechanical forces to intersect with other biochemical signaling pathways rather than acting as isolated inputs. The LATS1/2-YAP axis integrates these signals (cytoskeletal tension, calcium, MAPK activity, and growth factor pathways) into transcriptional programs governing proliferation, survival, and ECM remodeling [42, 43].

TGF*β* plays a central role across this network. Stretch activates it through cleavage of latent TGF*β* stored in the ECM [44], and an autocrine feedback loop sustains its signaling [45]. It propagates through both canonical Smad pathways [46, 47] and noncanonical axes including ERK [48], PI3K [49], and sGC/cGMP/PKG [50], coordinating transcriptional responses involved in cell adhesion and ECM remodeling.

To reflect the diverse MMP roles, MMP1 was encoded as a direct collagen degrader (inhibiting CI and CIII), MMP2 as both a collagen degrader and an activator of latent TGF*β*, and MMP9 as an activator of latent TGF*β* only.

### Model calibration

We used two types of experimental datasets from existing literature for model calibration: (1) input stimulation data and (2) node silencing data shown in Fig 2. The first dataset is built from studies that stimulate the corresponding model inputs and measure the expression/activation of downstream nodes. For this, we gathered a total of 92 data points across two complementary datasets: 62 from input-stimulation experiments and 30 from knockdown experiments. The input-stimulation dataset is distributed by stimulus as follows: 18/62 mechanical, 13/62 TGF*β* infusion, 9/62 AngII, 1/62 IL6, 8/62 FGF, 5/62 PDGF, 4/62 ET1, 3/62 TNF*α*, and 1/62 IL1. The knockdown dataset is distributed by silenced node as follows: 12/30 TGF*β* infusion, 7/30 IL6, 6/30 TNF*α*, and 5/30 SMAD3. For mechanical stimulus input, we carefully selected in vitro studies that defined baseline as cultures in substrates with edges restricted and no stretch applied. The corresponding stimulated state is 20%-40% static strain. Also, the experimental node activation was measured after 1 hr to 7 days of stretching. For infusion of TGF*β* and other biochemical nodes, stimulation was usually done through media supplementation with recombinant species. Cell cultures were performed using human dermal fibroblasts or equivalent mouse cell lines. For the second dataset, we collected node-silencing data from studies with gene KO, siRNA treatments, or protein inhibitors. Although they have associated different levels of silencing effectiveness, the silenced node is modeled with *Y*_*max*_ = 0 (assumed to be 100% effective).

**Figure 2:**
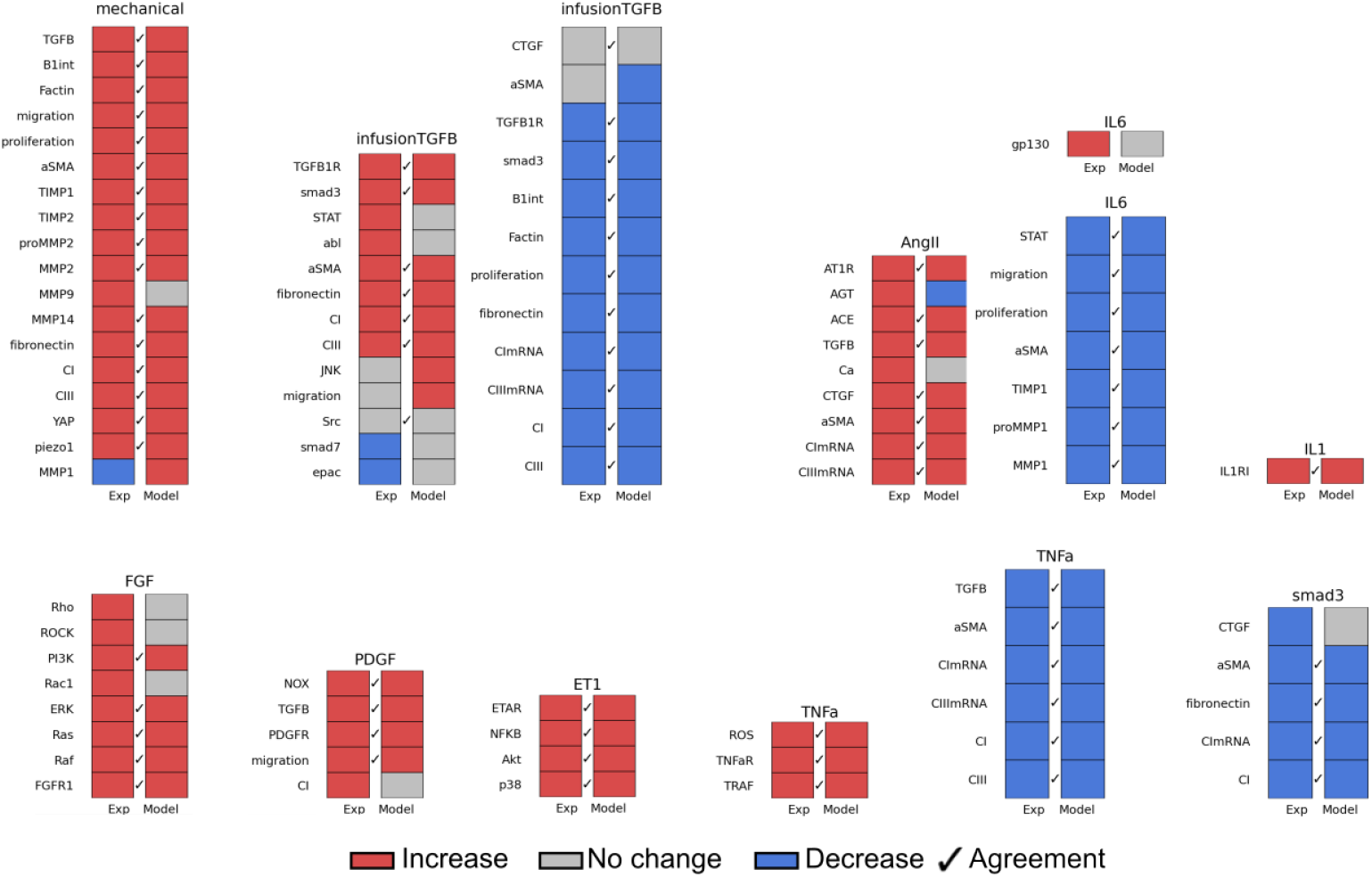
Agreement between network model predictions and experimental observations across a set of signaling perturbations. Each panel corresponds to a specific input stimulation or node inhibition (KO). Red, gray, and blue boxes denote increased, unchanged, and decreased activity, respectively. Check marks indicate qualitative agreement between model predictions and experimental observations.

We performed a two-step calibration using the Blackbox package in Python [51], considering two calibration datasets through the objective function defined in Suppl. 2. The first step was to calibrate the initial 4 parameters: *EC*50, *n, b*, and *p. EC*50 and *n* correspond to the half-maximal activation and the Hill coefficient for all interactions in the network. *b* and *p* are defined to be the baseline and perturbation values of the inputs such that (see Suppl. 2 for more details):

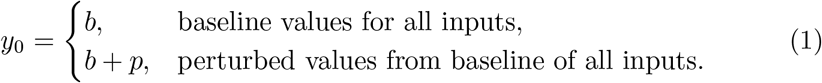

These 4 parameters were then fixed and we proceeded to fine tune the corresponding reaction weights **w**, which had been set to one for all reactions during the calibration of *EC*50, *n, b*, and *p*. The vector of reaction weights, **w**, however, has too many degrees of freedom (one per reaction). To increase interpretability and regularize the optimization, reactions were initially clustered to reduce dimensionality using k-means clustering, and get the number of clusters following elbow method. We calibrated the weight of each cluster reaction using the same optimization scheme used for *EC*50, *n, b*, and *p*.

### Full knock-out (KO) sensitivity analysis varying tension levels

We conducted a full KO sensitivity analysis of the system, in which we silenced one node at a time and recorded the steady-state activity changes in the other nodes. This analysis helps to understand how influential and sensitive the nodes are relative to the activity of other nodes, since it reveals nodes that serve as control points, propagating signals broadly, or that are highly sensitive to upstream nodes. To analyze the effect of each node removal in the network, we analyze changes in the steady state after performing a full KO by setting *Y*_*max*_ from 1 to 0. From the full KO, we define *influence* of node Y on node X as the change in activity (Δ_*Y*_ *X*) that node Y KO causes on node X. We performed the full KO of the system at different tension levels:

- Low tension (*y*_*mechanical*_ = 0.01)
- Baseline tension (*y*_*mechanical*_ = *b*)
- High tension (*y*_*mechanical*_ = *b* + *p*)
- Maximum tension (*y*_*mechanical*_ = 1)

### Coupling between signaling framework and one-dimensional growth

We considered uniaxial stretching of skin tissue and calibrated the model using a published dataset [9], in which directional total stretch *λ*(*t*), prestretch *λ*_*p*_(*t*), and growth stretch *λ*_*g*_(*t*) were reported, specifically along the caudal-rostral direction. Growth is represented by a scalar growth stretch *λ*_*g*_(*t*), such that the 1D multiplicative decomposition reduces to

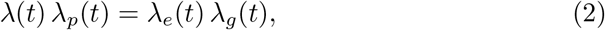

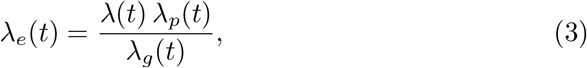

where *λ*_*e*_(*t*) is the elastic stretch driving mechanotransduction. Mechanical stimulation to the fibroblast signaling network was defined as a linear function of elastic stretch,

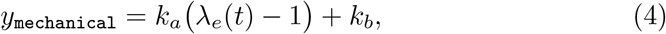

and used to set the model mechanical input (node `mechanical` of the ODE signaling system) at each time step. The signaling dynamics were advanced with the standard logic-based ODE system, yielding the instantaneous collagen-I production rate (node `CI`) with rate *ĊI*(*t*).

Collagen production was then mapped to a nonnegative growth stimulus via

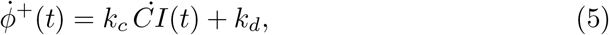

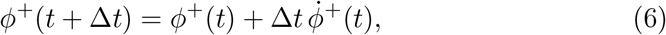

and the scalar growth stretch evolved according to

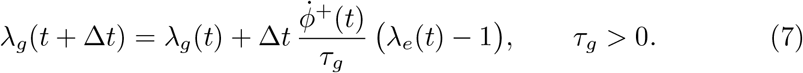

The parameters *k*_*a*_ and *k*_*b*_ (mechanical-to-signal mapping), *k*_*c*_ and *k*_*d*_ (collagen-to-growth mapping), and *τ*_*g*_ (growth time constant) were calibrated using the experimental dataset by minimizing the discrepancy between predicted and measured *λ*_*g*_(*t*) trajectories across all reported time points.

### Anisotropic growth tensor

To extend the calibrated one-dimensional coupling to directional growth, i.e. the estimation of the full growth tensor **F**_*g*_, we replaced the scalar growth variable by an orientation-dependent growth stretch *λ*_*g*_(*θ, t*), where *θ* ∈ [0, *π*) denotes an in-plane material direction. The angular domain was discretized into *N*_*θ*_ equally spaced orientations, each represented by the unit vector

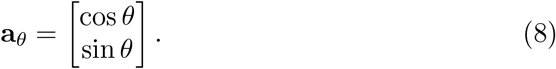

For each direction, the directional total stretch was computed from the prescribed deformation through the right Cauchy-Green tensor,

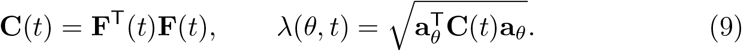

In the present implementation, the applied deformation was assumed to follow strip-biaxial kinematics,

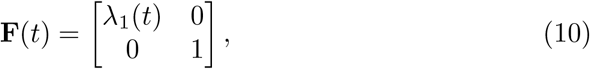

while the prestretch was assumed isotropic and trivially set to 1. Note that the strip-biaxial deformation **F** is arbitrary in this example, it serves as a way to illustrate the extension of the 1D calibration to the tensorial setting but the formulation works for arbitrary **F**. In other words, once the one-dimensional growth model is calibrated, the extension to a growth tensor is simply the result of integrating contributions along the unit circle.

The elastic stretch in each direction is computed from the multiplicative decomposition as

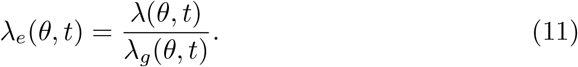

Unlike the one-dimensional setup, in which signaling was evaluated only along the loading direction, the directional extension assigns an independent signaling state to each angular direction. For each *θ*, the local elastic stretch *λ*_*e*_(*θ, t*) is mapped to a mechanical input for the signaling network, yielding an orientation-specific signaling response and collagen production rate 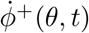.

Growth is then updated independently in each direction according to

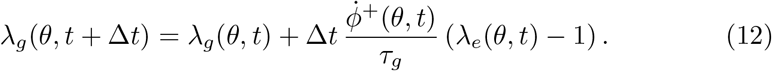

This formulation allows anisotropic growth to emerge from the coupling of kinematics driven by **F** and the direction-specific signaling response.

## Results

### Input-output relations

We aimed to identify the parameter combination that best fits the data. We assessed the percentage of qualitative agreement with experimental results following input stimulus and node KO, compared to baseline conditions. We performed an initial fit over four global parameters (*EC*50, *n, b*, and *p*), as described in the Methods Section. With this initial fitting, the model accurately predicted 65% of the collected data. To further improve the accuracy, we fixed these four parameters after the initial optimization and proceeded to optimize the interaction weights, as described in the Methods Section.

The weights were clustered into five families via the elbow method to determine the optimal number of clusters (Fig. S1). The five interaction families selected with this method have clear biological interpretation and correspond to *ECM turnover, Mechanosensitive contractile program, Inflammatory stress & immune remodeling, Mechanotransduction & cytoskeleton remodeling*, and *TGFβ-MAPK-MMPs activation* (Fig. 3).

**Figure 3:**
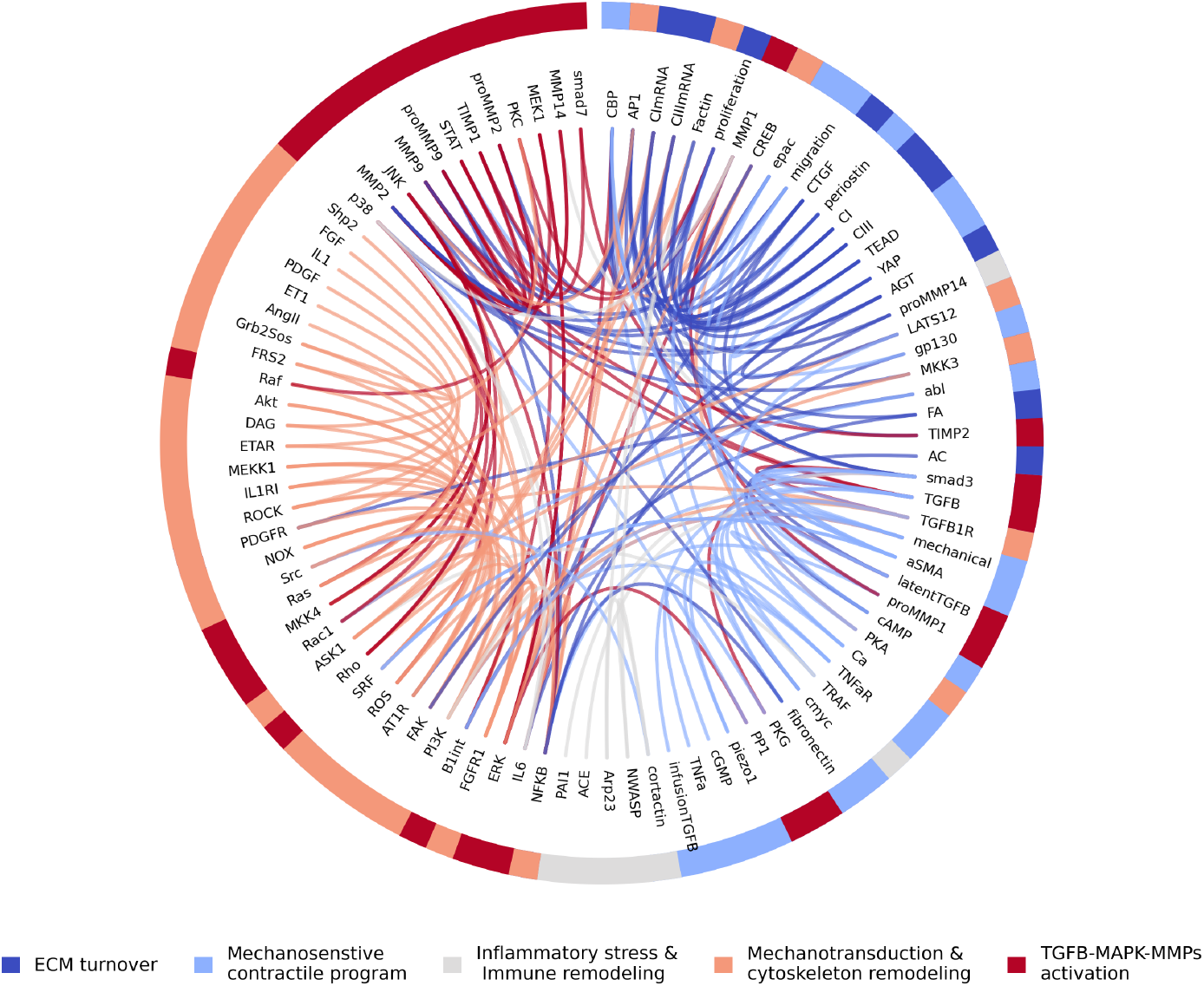
Clustering analysis of the fibroblast signaling network organized by biochemical reaction families. Nodes are arranged along the circle and grouped into five functional families. Colored outer arcs indicate family membership, while chords represent regulatory interactions between nodes. The density and distribution of chords highlight the dominant inter-family communication patterns that govern fibroblast activation and matrix remodeling in the model.

The *ECM turnover* family captures collagen synthesis, matrix degradation (MMPs/TIMPs), and transcriptional regulation of ECM-associated genes including fibronectin and CTGF. The *Mechanosensitive contractile program* integrates mechanical sensing (e.g., PIEZO1/Ca^2+^ and YAP signaling) with SMAD/SRF-mediated transcription to drive *α*SMA expression and myofibroblast activation. The *Inflammatory stress & immune remodeling* family encompasses p38-, NF*κ*B-, and cytokine-associated signaling, including actin regulators that modulate inflammatory transcriptional responses. The *Mechanotransduction & cytoskeleton remodeling* family represents integrin-FAK-Rho/ROCK-actin pathways that couple extracellular mechanical cues to intracellular force transmission and MAPK activation. Finally, the *TGFβ-MAPK-MMPs activation* family captures the regulatory axis linking TGF*β* activation, MAPK signaling, and protease-mediated matrix remodeling.

Further optimization of the five clustered interaction weights, after fixing *EC*50, *n, b*, and *p*, reproduced 75/92 of the curated experimental responses (82% overall agreement) on the training set. The set of calibrated parameter values is reported in Table 1. When stratified by validation dataset, the model reproduced 28/30 KO responses (93%) and 47/62 stimulation responses (76%). The high KO accuracy confirms that the core regulatory architecture is robust, particularly for perturbations affecting TGF*β* signaling and mechanically driven pathways. Under stimulation conditions, the model successfully captured the majority of mechanical- and TGF*β*-induced responses, including consistent trends in collagen production (CI, CIII), fibronectin, CTGF, and proliferation-related outputs across time points.

**Table 1:** Calibrated parameters of the dermal fibroblast signaling model and tissue-cell coupling framework.

| Symbol | Description | Value | Units |
| --- | --- | --- | --- |
| $EC_{50}$ | Half-max activation threshold | 0.59 | [-] |
| $n$ | Hill coefficient | 1.40 | [-] |
| $b$ | Baseline input level | 0.28 | [-] |
| $p$ | Input perturbation magnitude | 0.65 | [-] |
| $w_1$ | ECM turnover weight | 1.02 | [-] |
| $w_2$ | Contractile program weight | 1.10 | [-] |
| $w_3$ | Inflammatory remodeling weight | 0.95 | [-] |
| $w_4$ | Cytoskeleton remodeling weight | 1.10 | [-] |
| $w_5$ | TGFβ-MAPK-MMP weight | 0.63 | [-] |
| $k_a$ | Mechanical-input slope | -98.12 | [-] |
| $k_b$ | Mechanical-input offset | -17.47 | [-] |
| $k_c$ | Collagen-to-stimulus gain | 82.09 | [-] |
| $k_d$ | Stimulus baseline offset | 1.24 | [1/h] |
| $\tau_g$ | Growth time constant | 94.69 | [-] |

### Tension-dependent shifts in node influence across fibroblast phenotypes

Fibroblast signaling and ECM remodeling pathways have been mapped extensively, yet the relative influence of individual regulators within these pathways remains poorly understood, especially under varying mechanical tension. To address this gap, we quantified how each upstream species and input perturbation shapes phenotypic outputs across four tension levels, defined as low (*y*_mechanical_ = 0.01), baseline (*y*_mechanical_ = *b*), high (*y*_mechanical_ = *b* + *p*), and maximum (*y*_mechanical_ = 1). For each tension level we identify “brakes” and “boosters”. “Brakes” are nodes *Y* with positive *influence* on a particular node X: Δ_*Y*_ *X >* 0. Meaning that under normal conditions, they downregulate node X. “Boosters” are nodes *Y* with negative *influence* on a particular node X: Δ_*Y*_ *X <* 0. That is, they normally upregulate node X. We quantified their effect on eleven phenotypic nodes (*α*SMA, proMMP1, proMMP2, proMMP9, MMP1, MMP2, MMP9, CImRNA, CI, fibronectin, and proliferation; Fig. 4). On each panel of Fig. 4A-D, the columns represent the union of the top-10 most influential KO identified for each phenotype within each tension context (eleven phenotypes *×* four tensions = 44 lists), extended from Fig. S2. Columns are ordered left-to-right by how frequently each node appears across these lists. When two nodes appear in the same number of lists, the one with the larger average *influence* across phenotypes and tensions is placed to the left.

**Figure 4:**
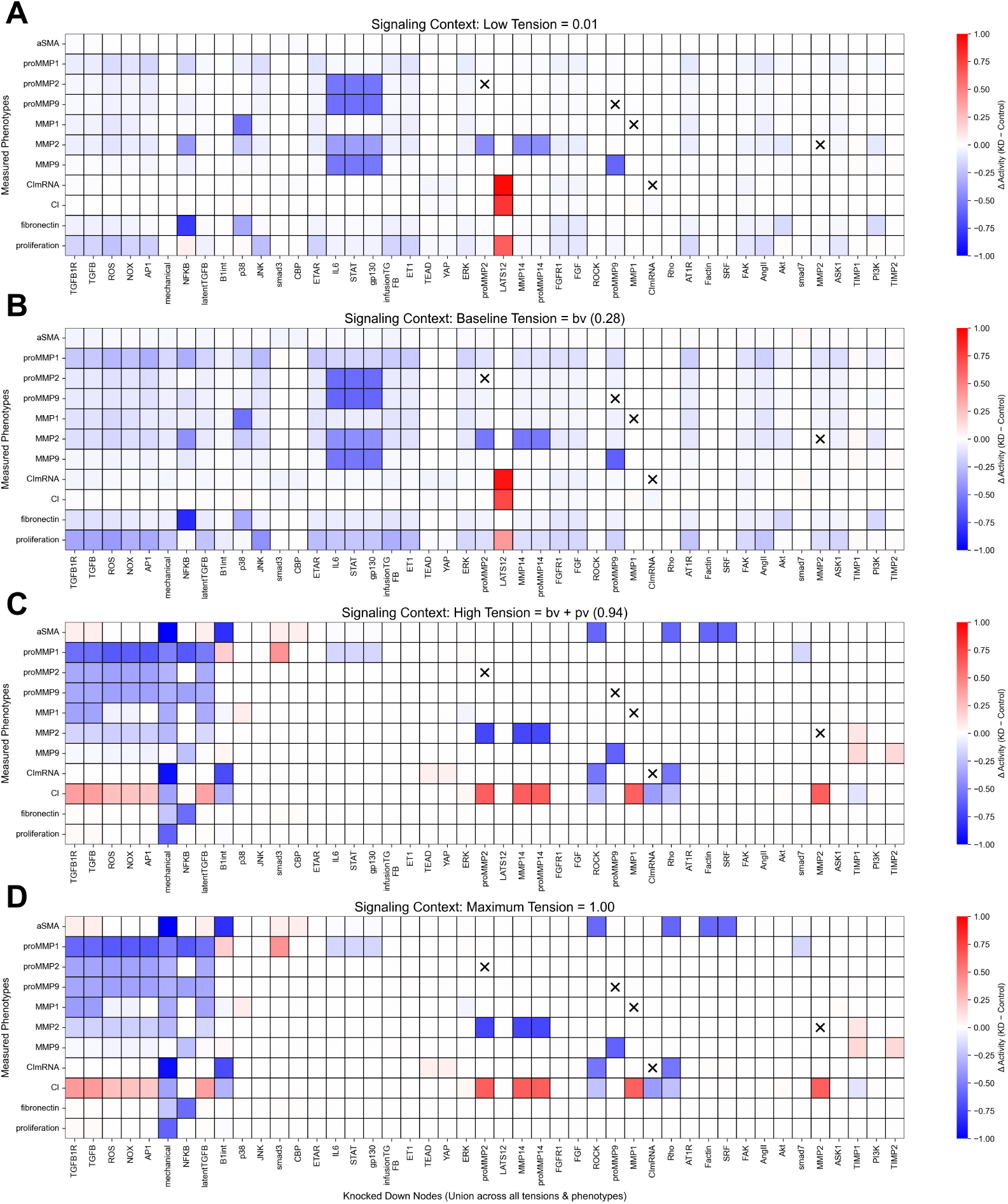
Tension-dependent regulatory influence across fibroblast phenotypic outputs. Each panel shows the change in steady-state activity of phenotypic nodes (rows) upon individual knockout of upstream nodes (columns) at four tension levels: low (A), baseline (B), high (C), and maximum (D). Positive values (red) indicate that removal of a node increases output activity (brake), while negative values (blue) indicate that removal decreases output activity (booster). “X” marks indicate grids where a phenotypical node is matched with its own KO. Columns are ordered left-to-right by how frequently each node appears across these lists. When two nodes appear in the same number of lists, the one with the larger average *influence* is placed to the left.

At low and baseline tension levels, regulatory influence is mostly activation-driven (Fig. 4A-B). (pro)MMP1/2/9 and fibronectin are controlled by distributed booster modules; only the collagen-related nodes (CI, CImRNA) and proliferation show a consistent brake, and that brake is a single node, LATS1/2 (Δ_LATS1/2_CI = +0.72, Δ_LATS1/2_CImRNA = +0.93, and Δ_LATS1/2_proliferation = +0.40). Several signaling modules act as broad boosters across multiple phenotypic nodes. Input nodes (TGF*β*, ET1, AngII, mechanical) and intermediate nodes (ROS, AP1, NOX, ERK) collectively produce a distributed activation of MMP1/2, proMMP1/2/9, fibronectin, and proliferation, with ΔX = − 0.14 ± 0.08 across these regulator-target combinations. *α*SMA is the least regulated phenotype at baseline; no upstream node produces Δ*α*SMA above 0.04.

Taken together, baseline tension defines an activation-dominated regime in which most phenotypic outputs are governed by combined booster modules with similar ΔX ranges, with consistent braking control coming only from LATS1/2 on collagen-related nodes and proliferation.

At high tension, regulatory control becomes more pronounced and structured (Fig. 4C). Boosters remain the dominant regulators, but several previously inactive brakes now become visible specifically in *α*SMA, proMMP1, MMP1, MMP2, MMP9, and CImRNA, which now have both boosters and brakes among their top influencers (Fig. S2C). The remaining phenotypes (proMMP2, proMMP9, fibronectin, and proliferation) instead show fewer but stronger boosters.

*α*SMA shifts from being weakly regulated at baseline to being strongly regulated at high tension by nodes along the integrin-Rho-ROCK-F-actin-SRF axis (Δ*α*SMA = − 0.64 ± 0.09). Its dominant brakes are TGF*β*R and SMAD3 (Δ*α*SMA = +0.07 ± 0.00).

Regarding ECM intermediate regulators, proMMP1 retains the same set of distributed boosters from baseline (TGF*β*, ROS, NOX, AP1, mechanical, NF*κ*B), but their mean influence amplifies from ΔproMMP1 = − 0.25 ± 0.06 at baseline to − 0.61 ± 0.06 at high tension. On the brake side, Smad3 emerges as a proMMP1 brake at high tension, having been essentially inactive at baseline (Δ_Smad3_proMMP1 = +0.01 at baseline, +0.43 at high tension). For MMP1, high tension sees TGF*β* family members and the mechanical input as the new dominant boosters (ΔMMP1 = −0.27 ± 0.15). Notably, p38 reverses its role on MMP1, shifting from the dominant MMP1 booster at baseline (Δ_p38_MMP1 = − 0.55) to a modest MMP1 brake at high tension (Δ_p38_MMP1 = +0.07). For MMP2, the ROS, NOX, and TGF*β* nodes gain booster influence (ΔMMP2 = −0.18±0.01 at high tension,while TIMP1 strengthens as a brake, Δ_TIMP1_MMP2 = +0.02 at baseline to Δ_TIMP1_MMP2 = +0.09 at high tension. MMP9 follows a similar redistribution, its dominant booster shifts from IL6 at baseline to NF*κ*B at high tension (Δ_NF*κ*B_MMP9 = −0.24), and TIMP1/2 strengthen as brakes (Δ_TIMP1/2_MMP9 = +0.15 ± 0.00).

Collagen shows control redistribution at both transcription and protein level. At baseline, dominant brake of CImRNA was LATS1/2. At high tension, YAP-TEAD takes over as a modest brake (Δ_YAP-TEAD_CImRNA = +0.06 ± 0.00). The mechanotransduction axis (integrin, Rho, ROCK) becomes the main booster (ΔCImRNA = − 0.59 ± 0.08). At the protein level, collagen (CI) becomes brake-dominated. Its main brakes are MMP1, MMP2, and MMP14 (ΔCI = +0.62 ± 0.00), with additional braking input from TGF*β*R, ROS, AP1, and NOX (ΔCI = +0.27 ± 0.07). Booster effects on CI remain weak: CImRNA, the integrin-Rho-ROCK axis, and TIMP1 (ΔCI = − 0.26 ± 0.09). Fibronectin and proliferation see pronounced control consolidation at high tension, with mechanical input the most dominant booster.

These shifts are accompanied by the emergence of two regulators that were nearly silent at baseline. Integrin (ITGB1) had a maximum | Δ_ITGB1_*X* | of 0.07 across all eleven phenotyping nodes *X* at baseline; at high tension this rises to |Δ_ITGB1_*X* | = 0.82, with strong effects on *α*SMA (− 0.82, booster) and proMMP1 (+0.19, brake). The mechanical input itself shows the same pattern where it becomes the dominant booster for fibronectin and proliferation as noted above. At maximum tension, the regulatory architecture is essentially saturated.

The mean change in |ΔX| from high to maximum tension across all eleven phenotypic nodes is small (+0.0002 ± 0.002) (Fig. 4D).

### TGFβ-mechanical crosstalk

We want to understand crosstalk between TGF*β* and mechanical signals since both are known regulators of dermal fibroblast activation [52, 53, 54]. To this end, we study dose-response profiles of relevant ECM remodeling outputs following TGF*β* stimulation at different tension levels (Fig. 5).

**Figure 5:**
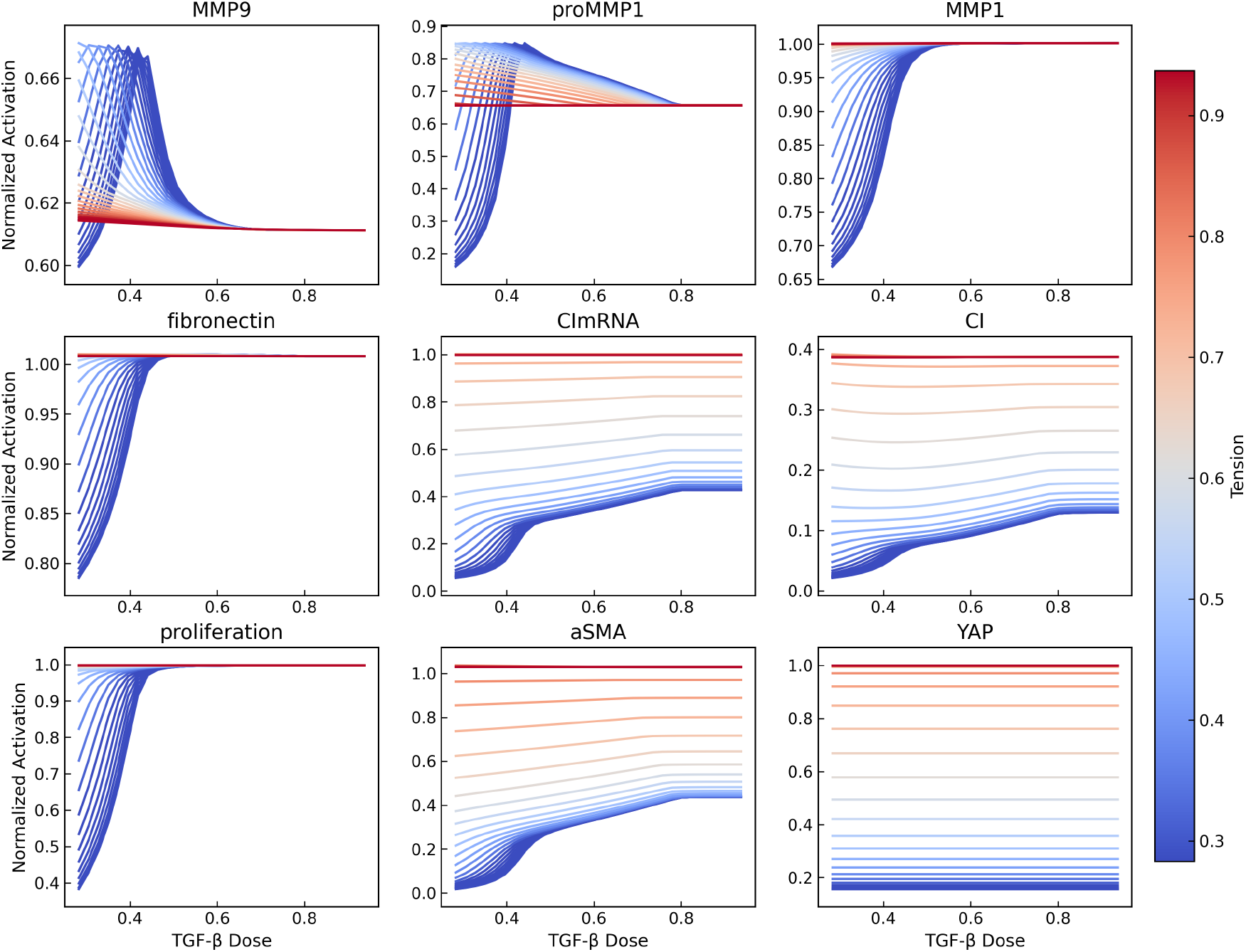
Dose-response behavior of fibroblast phenotypic nodes under combined TGF*β* stimulation and mechanical tension. Each panel shows the normalized activation of a model species as a function of TGF*β* dose, with color indicating tension level.

Fig. 5, top row, focuses on ECM degrading enzymes and their gene expression. MMP9 shows a peak at intermediate TGF*β* doses, then drops slightly and settles at a lower steady value as TGF*β* increases further. When tension increases, the peak shifts to the left, meaning the onset shifts to lower TGF*β* doses, increasing its sensitivity to TGF*β*, and eventually very high tension saturates MMP9 response to TGF*β*. proMMP1 displays a similar non-monotonic response to TGF*β* that depends strongly on the tension context. The non-monotonic behavior suggests an optimal stimulation range rather than a purely dose-dependent increase at low tension. As tension increases, a shift of the peak toward lower TGF*β* suggests increased sensitivity.At the highest tension levels, mechanical input alone brings proMMP1 near saturation, and TGF*β* has little room to further modulate the response. MMP1, in contrast, exhibits monotonic increases in activation with increasing TGF*β* dose, reaching saturation at approximately half of the maximal input. Increasing tension shifts the dose–response curves toward lower TGF*β* doses, effectively increasing sensitivity to TGF*β*.

Fig. 5, middle row, focuses on ECM biopolymers. Fibronectin displays a dose–response behavior similar to that observed for MMP1, with a monotonic increase and early saturation at low tension. As tension increases, the dose-response curve shifts toward lower TGF*β* doses, increasing sensitivity to TGF*β*; at the highest tensions, mechanical input alone brings fibronectin near saturation, so the response converges to a high activation state across all doses. CImRNA and CI show monotonically increasing dose-response with respect to both TGF*β* and tension. Although CImRNA and CI exhibit broadly similar TGF*β* dose–response behavior, their ranges are different. The gap between the CImRNA expression and the actual CI accumulation shows the role of TIMP1, MMP1, and MMP2 in limiting collagen accumulation even as transcription increases [55].

The third row in Fig. 5 focuses on cell phenotype. Proliferation increases with TGF*β* dose at low tension levels, reaching saturation quickly with respect to TGF*β* concentration. As tension increases, proliferation becomes elevated even at very low TGF*β* doses reflecting a leftward shift of the dose-response curve and increased sensitivity to TGF*β*. At high tension, mechanical input alone brings proliferation near its steady-state maximum, producing a nearly flat dose-response curve. Fibroblast contractility, captured by *α*SMA activation, shows the same trends as CImRNA and CI. Finally, our model shows that YAP activation is strongly tension-dependent but remains essentially unchanged across the entire TGF*β* dose range.

### Model validation against additional experimental data

Fig. 6A compares experimental measurements of TGF*β*–driven fibrotic signaling in a bleomycin-induced mouse model of skin fibrosis with corresponding model predictions. Experimentally, bleomycin treatment (BLEO) induces a marked increase in SMAD2/3 phosphorylation, *α*SMA expression, and dermal thickness relative to controls. This aligns with activation of canonical TGF*β* signaling and myofibroblast differentiation [56]. Vehicle-treated animals (VEHIC) exhibit responses comparable to those of BLEO, indicating that the vehicle alone has no protective effect. In contrast, TGF*β* pathway inhibition (aTGF) significantly suppresses p-SMAD2/3 levels, reduces *α*SMA–positive cells, and attenuates dermal thickening. Model simulations reproduce these qualitative trends: baseline conditions show minimal signaling activity, induction of TGF*β* signaling recapitulates the profibrotic BLEO/VEHIC phenotype, and simulated TGF*β* receptor blockade effectively reduces SMAD activation and downstream fibrotic outputs. These results demonstrate that the model captures the dominant role of TGF*β*/SMAD signaling in bleomycin-induced dermal fibrosis and its reversal by pathway inhibition, in agreement with in vivo observations.

**Figure 6:**
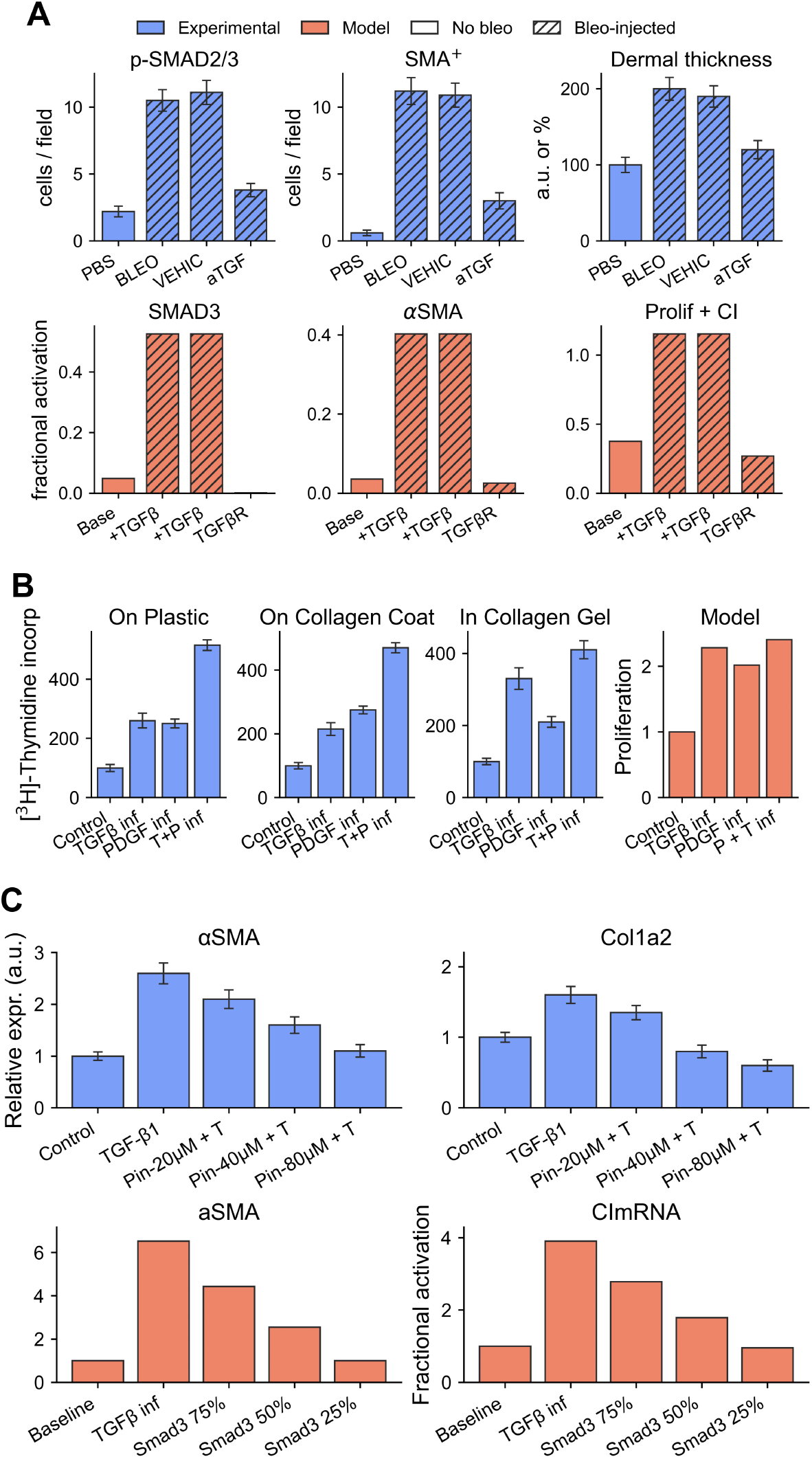
Comparison between experimental observations and model predictions across multiple fibrotic signaling contexts. (A) p-SMAD2/3^+^ and *α*SMA^+^ cell counts and dermal thickness under PBS, bleomycin (BLEO), BLEO+vehicle (VEHIC), and BLEO+anti-TGF*β* (aTGF) [56], vs. model SMAD3, *α*SMA, and combined proliferation+collagen I outputs. (B) Fibroblast proliferation ([^3^H]-thymidine incorporation) under control, TGF*β*, PDGF, and TGF*β*+PDGF stimulation on plastic, collagen-coat, and collagen-gel substrates [57], vs. model proliferation predictions. (C) Relative *α*SMA and Col1a2 expression under TGF*β ±* pinocembrin (TGF*β*-SMAD3 inhibitor) [58], vs. model *α*SMA/CImRNA activation under simulated graded SMAD3 inhibition. 18

Fig. 6B summarizes data from dermal fibroblasts cultured on plastic, collagen-coated surfaces, or embedded within collagen gels and stimulated with PDGF and TGF*β* infusion [57]. Thymidine incorporation reflects DNA synthesis and cell proliferation activity. Among the three experimental conditions, fibroblasts cultured within collagen gels represent the environment most comparable to native skin and the context of the datasets used to construct our signaling network. Fig. 6B, (right panel) most closely follow the experimental responses observed in collagen-gel cultures, correctly capturing the relative increase in proliferation induced by PDGF and TGF*β* under baseline tension.

Fig. 6C shows experimental results that quantified the expression of *α*-SMA and Col1a2 following TGF*β* stimulation alone or in combination with pinocembrin at multiple doses [58]. Pinocembrin is a known inhibitor of the TGF*β*-SMAD3 signaling pathway. To reproduce these conditions in our model, we simulated (i) baseline, (ii) TGF*β* infusion by increasing the TGF*β* input node, and (iii) TGF*β* + pinocembrin by reducing the maximum activation parameter *Y*_max_ of the SMAD3 node to mimic graded inhibition (75%, 50%, and 25%). The model shows the qualitative trends reported experimentally: TGF*β* increases profibrotic markers, while progressive SMAD3 inhibition attenuates these responses.

### Prediction of skin growth from tissue-cell coupled feedback

Fig. 7 summarizes the validation of the coupled tissue-cell framework and its extension to anisotropic growth predictions and Table 1 lists values of coupling parameters. Fig.7 A compares predicted and experimental growth stretch *λ*_*g*_ at 24, 72, and 168 hours. Across all time points, the model reproduces the overall increase in growth magnitude, with both predicted and measured values shifting toward larger *λ*_*g*_ at later times. The scatter plots show a positive correlation between model predictions and experimental measurements, indicating that the coupled mechanobiological formulation captures the general trend and order of magnitude of directional growth along the loading direction. Fig.7 B shows the aggregated comparison across all samples and time points, where model predictions remain distributed within the experimental range despite the larger dispersion observed at later times.

**Figure 7:**
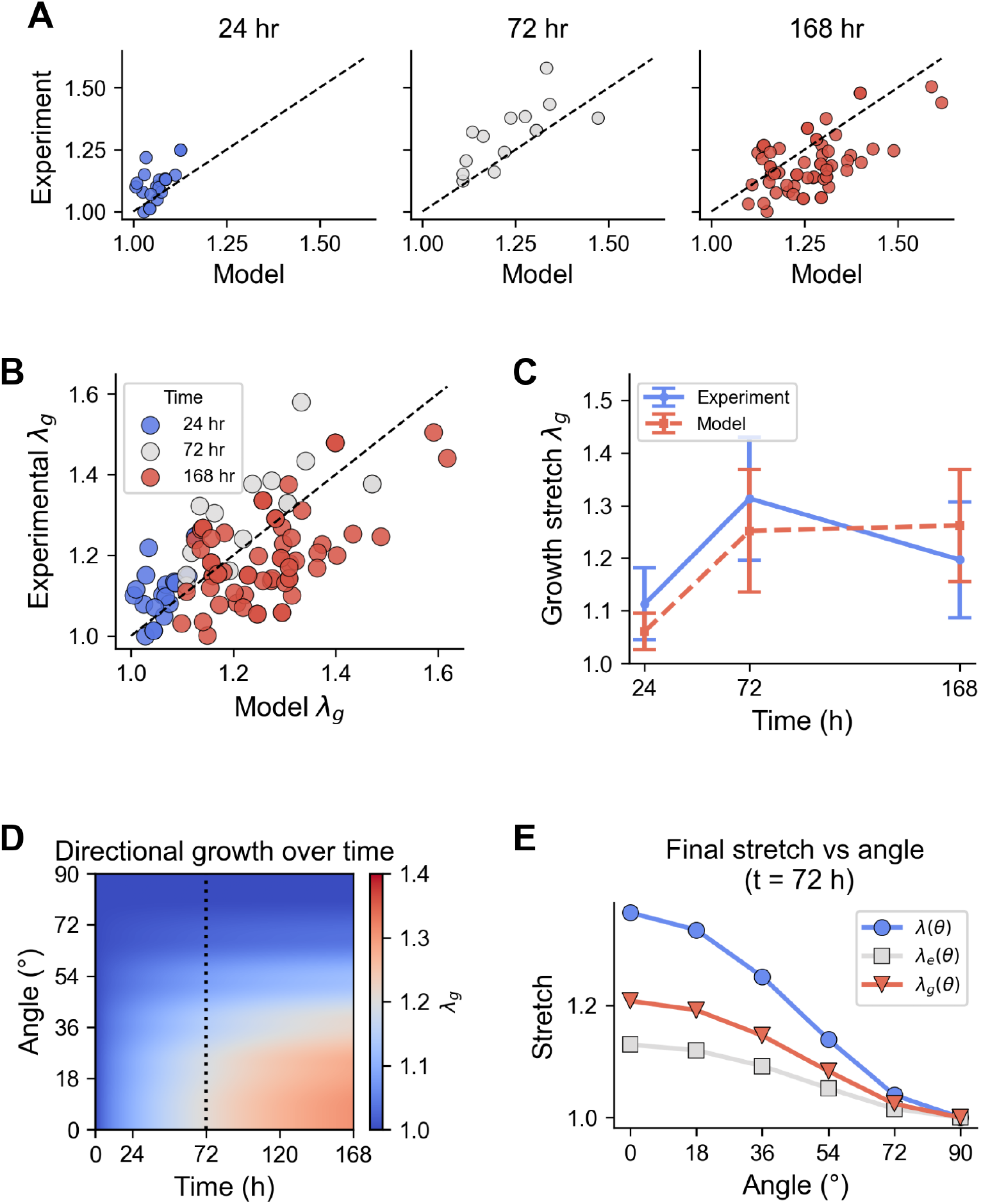
Validation of the tissue-cell coupling framework and prediction of anisotropic growth. (A) Comparison between model predictions and experimental measurements of growth stretch *λ*_*g*_ along the loading direction at 24, 72, and 168 h. (B) Combined scatter comparison across all samples and time points. (C) Mean and standard deviation of experimental and predicted growth stretch over time. (D) Predicted evolution of directional growth *λ*_*g*_ (*θ, t*). The vertical dotted line marks *t* = 72 h, the time point used for the angular decomposition shown in panel E. (E) Angular decomposition of total stretch *λ*(*θ*), elastic stretch *λ*_*e*_(*θ*), and growth stretch *λ*_*g*_ (*θ*) at *t* = 72 h for the same sample as in panel D.

Fig.7 C summarizes the temporal evolution of growth stretch by comparing the mean and standard deviation of experimental measurements with model predictions. The model captures the increase in growth from 24 to 72 hours and remains within the experimental variability at later times, demonstrating that the calibrated framework reproduces the overall time-dependent growth response under sustained loading.

To investigate how the calibrated coupling framework extends to anisotropic growth behavior, the scalar growth stretch *λ*_*g*_(*t*) was generalized to an orientation-dependent quantity *λ*_*g*_(*θ, t*) by discretizing in-plane material directions. For each orientation *θ*, the total directional stretch *λ*(*θ, t*) was obtained from a prescribed deformation tensor, and the corresponding elastic stretch *λ*_*e*_(*θ, t*) depended on the growth along each particular direction, *λ*_*g*_(*θ, t*).

Fig. 7 D illustrates the predicted evolution of directional growth *λ*_*g*_(*θ, t*) over time. Growth increases predominantly near the loading direction (*θ* = 0^◦^) while remaining smaller toward transverse orientations, reflecting the anisotropic mechanical environment imposed by the deformation. The vertical dotted line indicates the time point considered for Fig. 7 E, which shows the corresponding angular profiles of total stretch *λ*(*θ*), elastic stretch *λ*_*e*_(*θ*), and growth stretch *λ*_*g*_(*θ*). The growth stretch is largest along the loading direction and decreases progressively toward transverse orientations.

Together, these results demonstrate that the calibrated tissue-cell coupling framework can reproduce experimentally observed growth in a porcine model of TE.

## Discussion

In this work, we developed a logic-based signaling model of dermal fibroblasts to investigate how biochemical cues and mechanical tension jointly regulate skin growth and remodeling during tissue expansion. The model builds on the logic-based ODE framework pioneered by Saucerman and colleagues for cardiac fibroblasts [24, 59, 39]. We adapted their network topology and modeling paradigm to the dermal context through systematic curation of mechanotransduction pathways, ligand inputs, and downstream phenotypic outputs relevant to skin G&R (see Supplemental materials and repository for a complete list of references for nodes and reactions). Extensive calibration against stimulation and KO datasets provides confidence in the biological fidelity of the resulting network (full list of references in Supplemental files and data repository).

One key finding is that the regulatory architecture of fibroblast signaling reshapes with mechanical tension (Fig. 4), shifting from a uniform, activation-driven regime at baseline to a narrow regulation regime at high tension in which a few nodes become extremely important. At low and baseline tension, the eleven phenotypic outputs we examined are mainly regulated by booster modules with similar effect magnitudes, with consistent inhibitory regulation coming only from LATS1/2 on collagen-related nodes and proliferation. At high tension, brakes become visible in the *α*SMA, proMMP1, MMP1, MMP2, MMP9, and CImRNA phenotypic nodes, which were substantially booster-controlled at baseline. Most of these brakes arise from initial boosters that change identity under high tension, most notably SMAD3 and p38. SMAD3 went from a soft booster to a strong proMMP1 brake, consistent with evidence from human dermal fibroblasts cultured on stiff plates [60] (data not used in the calibration stage). The role of p38 in regulating MMP-1 is dynamic in our modeling results, shifting from a strong promoter to a gentle inhibitor depending on context. This dynamic role has been found in contracting three-dimensional collagen lattices, where p38 promotes MMP-1 expression through NF-*κ*B activation while triggering counteracting mechanisms that restrict MMP-1 levels [61, 62].

High tension reveals the balance between Collagen-I transcription and protein regulation. At baseline, both CI and CImRNA are mainly inhibited by LATS1/2. Under high tension, CImRNA becomes activated by mechanotransduction (integrin-Rho-ROCK) with a reduction in YAP-TEAD braking, while CI protein is brake-dominated by MMPs action (MMP1/2/14), in agreement with experiments [42, 63, 64, 65].

Another finding is the regulation of TGF*β* signaling by mechanical tension (Fig. 5). MMP1, TIMP1, fibronectin, and proliferation each show a leftward shift of their dose-response curves as tension increases, meaning that tension increases sensitivity to TGF*β* signaling . This is consistent with additional experimental observations of strong fibronectin induction at low TGF*β* doses and saturation at higher doses [66], and with reported saturation of DNA synthesis in response to rising TGF*β* under low-tension conditions [67, 68]. The model also predicts a sigmoidal increase in *α*SMA with TGF*β* dose across all tension levels, in agreement with reported dose-dependent increases approaching a plateau at high TGF*β* [69]. Notably, YAP activation is predicted to be strongly tension-dependent while remaining insensitive to TGF*β* dose, consistent with independent experimental observations that TGF*β* does not directly increase YAP nuclear localization or alter YAP phosphorylation in human dermal fibroblasts [70].

Together, the calibration results, validation, and further consistency with literature not used during model development provide confidence that the network captures the dominant regulatory logic of dermal fibroblast signaling under tensile loading. With this model, we were uniquely positioned to study the multi-scale coupling of cellular signaling and tissue-level adaptation, such as skin growth in TE.

To model skin growth in response to stretch, we used the collagen production node from the fibroblast signaling network as the driver of the tissue-level growth stretch. The coupled model reproduced the magnitude and temporal progression of *λ*_*g*_ observed experimentally at 24, 72, and 168 h. Importantly, the coupling introduces a feedback loop that is absent from cell-level models alone: as the tissue grows, the elastic deformation (and therefore the mechanical input to the signaling network) also evolves over time. This bidirectional coupling distinguishes the present framework from prior work that has characterized signaling responses at fixed mechanical states [24, 25, 26], and is consistent in spirit with multiscale formulations in cardiac mechanics that integrate cell-level signaling with organ-level growth [27, 28].

### Model limitations and future directions

Several limitations of the present framework should be acknowledged. First, mechanical tension is represented as a kinematic input. In reality, ECM remodeling alters tissue stiffness, which in turn can change the strain experienced by cells. There is also direct coupling between stiffness and the mechanical input node of the signaling model which is not considered here [71].

Second, the model considers fibroblast signaling in isolation and does not account for interactions with other cell populations present in skin, including keratinocytes, endothelial cells, immune cells, and adipocytes. These populations can influence fibroblast behavior through paracrine signaling and ECM remodeling. Extending the framework to incorporate multicellular crosstalk would enable investigation of tissue-level coordination and inflammatory feed-back mechanisms relevant to other skin processes besides G&R, e.g. wound healing and fibrosis.

Third, the logic-based ODE framework emphasizes steady-state regulatory influence and qualitative trends rather than precise temporal dynamics. Transient responses, time delays, and oscillatory behaviors that may arise during rapid mechanotransductive or inflammatory signaling are not well captured by this formulation. Model parameters are calibrated to reproduce qualitative experimental trends rather than absolute quantitative measurements. That said, the coupling to tissue-level growth is quantitative, with only four transfer function parameters linking cell-level signaling outputs to tissue-level growth stretches. This architecture (qualitative at the cell level, quantitative at the tissue level) allows the model to leverage tissue-level mechanical data for quantitative calibration while still encoding the pathway-level logic needed to faithfully simulate the biological mechanism. Extending this framework to predict how pharmacological modulation of specific signaling nodes translates into changes in tissue-level growth rates remains an important direction for future work and would require additional cell-level experimental calibration.

## Conclusion

We developed and validated a multiscale model of fibroblast-driven skin growth and remodeling that integrates cell-level biochemical signaling with tissue-level growth. The signaling network was constructed through extensive literature curation, calibrated against stimulation and knockdown datasets, and validated across multiple independent experimental contexts. Coupling collagen production from the fibroblast signaling model to a tissue-level growth formulation reproduced experimentally measured growth dynamics during tissue expansion. By explicitly linking intracellular regulatory logic to tissue-level outcomes, the framework opens the door to mechanistic hypothesis testing and the in silico design of targeted interventions in tissue expansion. More broadly, the modeling approach is applicable to other settings in which skin growth and remodeling are governed by cell-level signaling, including wound healing and fibrosis.

## Author Contributions

**Laura Nuñez-Alvarez:** Conceptualization, Methodology, Software, Formal analysis, Investigation, Visualization, Writing. **Joanna K. Ledwon:** Data curation, Investigation. **Parul Rai:** Data curation. **Kathryn Reisner:** Data curation. **Tianhong Han:** Methodology, Software. **Luis Solorio:** Supervision, Funding acquisition, Resources. **Arun Gosain:** Supervision, Funding acquisition, Resources. **Adrian Buganza Tepole:** Conceptualization, Methodology, Formal analysis, Writing, Supervision, Funding acquisition.

## Acknowledgments

ABT acknowledges support from the National Institute of Arthritis and Musculoskeletal and Skin Diseases, National Institute of Health, United States under award R01AR074525.

## Supplementary Material

Code and data available at: https://github.com/lnunezal/TE_R01

## Notes

https://github.com/lnunezal/TE_R01

### Competing Interest Statement

The authors have declared no competing interest.

