## Supplemental material for "Coupling fibroblast mechanotransduction signaling to tissue growth in a multiscale model of skin expansion"

### Supplemental Information

#### 1 Network construction

The dermal fibroblast signaling network was adapted from a previously published cardiac fibroblast network. Nodes and interactions specific to the cardiovascular system were removed, nodes with documented evidence in dermal fibroblasts were retained, and three signaling axes relevant to skin growth and remodeling were added. The final network connects nine inputs to thirteen outputs through 96 nodes and 151 reactions. Each interaction was required to be supported by at least two independent experimental studies, prioritizing evidence in human dermal fibroblasts, mouse dermal fibroblasts, dermal fibroblast-like cells, and mammalian fibroblasts more broadly. Crosstalk interactions characterized in other cell types were retained when corroborated by at least one report in fibroblast-related cells. The full list of nodes, interactions, family assignments, and supporting references is provided in the spreadsheet `interaction_classification.xlsx` available in the code repository, organized into four sheets (node families, node-to-family assignment, interaction families, and interaction-to-family assignment).

#### 2 Model building using Netflux

The logical network was translated into a system of ordinary differential equations (ODEs) using Netflux, a publicly available logic-based ODE pipeline. Each node activity $y_i(t) \in [0, 1]$ is governed by

$$\tau_i \, \dot{y}_i(t) = f_i\big(\{y_j(t)\}_{j \in \mathcal{P}(i)}\big) \; y_{\max,i} - y_i(t), \tag{S1}$$

where $\mathcal{P}(i)$ denotes the set of upstream regulators of node $i$, $f_i$ is the continuous-valued logic function built from Hill activation/inhibition terms combined according to the network rule (AND, OR, NOT), $\tau_i$ is the node-specific time constant, and $y_{\max,i}$ is the maximum activation level. The exported ODE system, baseline parameter file, and integration utilities are included in the code repository as `NetfluxODE_5.py`, `NetfluxODE_5_params.py`, and `NetfluxODE_5_run.py`.

### 3 Parameterization

Model parameters are defined at the node and reaction levels. Let $N$ denote the number of nodes and $R$ the number of reactions in the network.

**Initial conditions.** The initial state of each node is defined as

$$y_{0,i} = \begin{cases} 0.1, & \text{non-input nodes under baseline conditions,} \\ b_i, & \text{input nodes under baseline conditions,} \\ b_i + p_i, & \text{input nodes under perturbed conditions,} \end{cases} \qquad i = 1, \ldots, N. \tag{S2}$$

**Maximum activation.** The maximum activation level is assumed to be identical for all nodes,

$$\mathbf{y}_{\max} = [1,\ 1,\ \ldots,\ 1]_N. \tag{S3}$$

**Time constants.** Reaction-specific time delays are prescribed as

$$\tau = \begin{cases} 0.1 \text{ h}, & \text{signaling reactions,} \\ 1 \text{ h}, & \text{transcription reactions,} \\ 10 \text{ h}, & \text{translation reactions.} \end{cases} \tag{S4}$$

**Reaction parameters.** Reaction weights, Hill coefficients, and activation thresholds were assumed uniform across reactions during the first stage of calibration,

$$\mathbf{w} = [1,\ 1,\ \ldots,\ 1]_R, \tag{S5}$$

$$\mathbf{n} = n_{\text{val}}\,[1,\ 1,\ \ldots,\ 1]_R, \tag{S6}$$

$$\mathbf{EC50} = \text{EC50}_{\text{val}}\,[1,\ 1,\ \ldots,\ 1]_R. \tag{S7}$$

The scalars $n_{\mathrm{val}}$ and $\mathrm{EC50}_{\mathrm{val}}$ were fit together with $b$ and $p$ in Stage 1; the per-reaction weights $\mathbf{w}$ were then refit in Stage 2 after reduction to five cluster weights (Section 4).

### 4 Model calibration

#### 4.1 Objective function

Model calibration was performed by minimizing a composite objective function defined as a weighted sum of mismatch penalties derived from two independent literature-curated datasets,

$$\mathcal{L}_{\mathrm{total}} = \alpha\,\mathcal{L}_{\mathrm{input}} + \beta\,\mathcal{L}_{\mathrm{KO}} + \mathcal{P}_B, \tag{S8}$$

where $\mathcal{L}_{\mathrm{input}}$ denotes the mismatch under input-stimulation experiments and $\mathcal{L}_{\mathrm{KO}}$ denotes the mismatch under knockout (KO) perturbations. Equal weights $\alpha = \beta = 1$ were used in all reported runs. $\mathcal{P}_B$ is a soft quadratic penalty introduced to keep activations bounded; it is non-zero when the dimensionless saturation $B = (EC50^n - 1)/(2EC50^n - 1) \leq 1$, with weight 10.

**Input stimulation mismatch.** $\mathcal{L}_{\mathrm{input}}$ is computed by simulating the model under perturbed input conditions and comparing the predicted change in steady-state activity, $\Delta y_{\mathrm{SS}} = y_{\mathrm{SS}}^{\mathrm{perturbed}} - y_{\mathrm{SS}}^{\mathrm{baseline}}$, with experimentally expected trends. Agreement is assessed using a ternary classification, where responses are categorized as increase, decrease, or no change relative to the unperturbed baseline.

**Knockout mismatch.** $\mathcal{L}_{\mathrm{KO}}$ is computed by simulating node-specific knockouts through modification of the maximum activation parameter ($y_{\max} = 0$ for the silenced node) and comparing the resulting $\Delta y_{\mathrm{SS}}$ values against expected knockout responses using the same ternary classification.

**Mismatch normalization.** Each mismatch term is normalized as the percentage of incorrect predictions,

$$\mathcal{L}_X = 100\,\% \;-\; \mathrm{accuracy}_X, \tag{S9}$$

such that larger values indicate poorer agreement between model predictions and expected behaviors. Predicted changes with $|\Delta y_{\mathrm{SS}}|$ below a prescribed threshold (0.05 for input stimulation, 0.02 for KO) are classified as no change.

#### 4.2 Two-stage optimization with Blackbox

Both calibration stages used the Blackbox global optimizer [1], a deterministic procedure that combines Latin-hypercube initial sampling with iterative refinement on a radial-basis-function surrogate model. Blackbox is well suited to expensive, low-to-moderate-dimensional objective functions in which each evaluation requires a complete steady-state simulation of the signaling network under both perturbed and baseline conditions.

**Stage 1.** The first stage optimized the four global scalars $EC_{50}$, $n$, $b$, and $p$, with all reaction weights fixed to unity. Search bounds were $EC_{50} \in (0.1, 1.0)$, $n \in (0.5, 3.0)$, $b \in (0.01, 0.5)$, and $p \in (0.01, 0.7)$, with the hard constraint $b + p \leq 1$. The optimizer converged on

$$EC_{50} = 0.59, \qquad n = 1.40, \qquad b = 0.28, \qquad p = 0.65,$$

with these values reproducing approximately 60/92 of the curated experimental responses (65% qualitative agreement). This first-stage fit captures the dominant Hill-saturation regime of the network but does not yet account for the per-reaction strength variation across signaling families, motivating a second optimization round.

**Stage 2.** The second stage held the four scalars fixed at their Stage 1 optima while the reaction weights were calibrated. The full vector of reaction weights $\mathbf{w}$ contains one entry per reaction; to regularize the optimization and improve interpretability, reactions were grouped into five biologically interpretable families via k-means clustering with the elbow criterion (Section 5), and a single weight per family was optimized. Search bounds were $w_i \in (0.01, 1.10)$ for each family.

The Stage 2 optimization was run with batch size 48 and an initial budget of 5,760 evaluations, extended across multiple invocations to 10,096 total evaluations. The best loss ($\mathcal{L}_{\text{total}} = 30.86$) was reached at evaluation 5,151 and remained unchanged across the subsequent $\sim$ 5,000 evaluations, indicating convergence under the Blackbox surrogate. The corresponding cluster weights are reported in Table 1 of the main text.

##### 4.3 From loss to qualitative agreement

The composite loss decomposes into per-dataset percent mismatches via Eq. (S9). At the best Stage 2 point, $\mathcal{L}_{\text{KO}} = 100 \times (1 - 28/30) = 6.67$ and $\mathcal{L}_{\text{input}} = 100 \times (1 - 47/62) = 24.19$, summing to $\mathcal{L}_{\text{total}} = 30.86$. We report two complementary accuracy metrics:

- the *pooled accuracy* across all 92 curated datapoints, $(28 + 47)/(30 + 62) = 75/92 = 82\%$, used in the main text;
- the *per-dataset average*, $(93.3 + 75.8)/2 = 84.6\%$, useful when KO and stimulation experiments are weighted equally regardless of dataset size.

The two differ because the stimulation dataset is roughly twice the size of the KO dataset; we use the pooled metric in the main text as it treats every literature-curated datapoint equally.

**Outlier exclusion.** No datapoints were excluded from the loss: all 30 KO responses (`KO_data_condensed2.xlsx`) and all 62 input-stimulation responses (`input_output_relations2.xlsx`) were retained throughout calibration.

**Reproducibility.** The optimization runner (`BB_run_DOF_cluster.py`), the objective function (`model_DOF_clustering_5.py`), and the full Stage 2 loss trajectory (`loss_log_DOF_cluster.csv`, 10,096 rows) are included in the code repository.

### 5 Reaction clustering

To reduce the dimensionality of the reaction-weight vector and regularize the Stage 2 optimization, the 151 reactions were grouped into families via k-means clustering on the binary node-incidence matrix (each reaction is a vector of 0/1 entries indicating which nodes participate). The optimal number of clusters was determined using the elbow method (Fig. S1), in which the within-cluster sum of squares (WCSS) is plotted against the number of clusters and the inflection point selected as the point beyond which further cluster reduction no longer significantly decreases the WCSS. Five clusters were selected. The resulting families have clear biological interpretation, corresponding to *ECM turnover*, *Mechanosensitive contractile program*, *Inflammatory stress & immune remodeling*, *Mechanotransduction & cytoskeleton remodeling*, and *TGFβ–MAPK–MMPs activation*, as described in the main text.

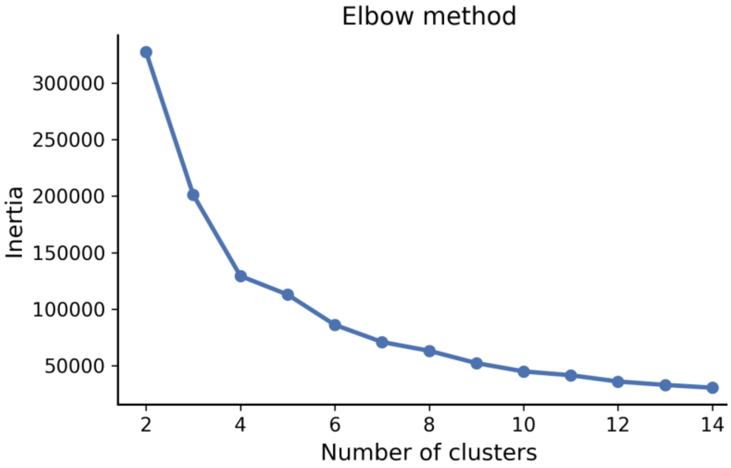


Figure S1: Elbow plot for clustering analysis. The y-axis represents the within-cluster sum of squares (WCSS) and the x-axis represents the number of clusters. Five clusters were selected as the inflection point at which further cluster-size reduction no longer significantly reduces WCSS.

### 6 Tissue-cell coupling calibration

The coupling between the calibrated signaling model and the tissue-level growth formulation introduces five additional parameters: $k_a$ and $k_b$ map the local elastic stretch to the mechanical input of the signaling network (Eq. 4 of the main text); $k_c$ and $k_d$ map the simulated collagen-I production rate to the growth stimulus (Eq. 5 of the main text); and $\tau_g$ is the growth time constant (Eq. 7 of the main text).

These five parameters were calibrated against the porcine tissue-expansion dataset [2], which reports directional total stretch $\lambda(t)$, prestretch $\lambda_p(t)$, and growth stretch $\lambda_g(t)$ at 24, 72, and 168 h. The objective function was the mean-squared error between predicted and measured $\lambda_g$ trajectories across all reported time points. Optimization was performed with Blackbox using bounds $k_a, k_c \in (-100, 100)$, $k_b, k_d \in (-100, 100)$, and $\tau_g \in (0.1, 100)$ h. The optimizer was run for 1,496 evaluations, with the best loss (3.87) reached at evaluation 1,361. The calibrated values are reported in Table 1 of the main text. The optimization runner (`coupling_G1_optimizer.py`) and the full loss trajectory (`optimization_log_G1_BB_all.csv`) are included in the code repository.

### 7 Per-phenotype regulatory analysis

Figure S2 presents the per-phenotype top-10 KO ranking that underlies the union-based view shown in Fig. 4 of the main text. For each tension context and each of the eleven phenotypic outputs analyzed, the ten knockdowns producing the largest absolute change in steady-state activity are shown, ranked left-to-right in decreasing order of influence. Self-knockdowns (i.e., the diagonal where the perturbed node equals the readout) are excluded. This complementary view exposes the per-phenotype regulator identities that are aggregated in the main-text union heatmap and provides the basis for the brake/booster classification used throughout the Tension-Dependent Shifts section.

### 8 Dose-response decomposition of non-monotonic responses

Two phenotypic outputs, MMP9 and proMMP1, display non-monotonic responses to TGF$\beta$ at low to intermediate mechanical tension. Figure S3 decomposes these responses into their upstream contributions to clarify the mechanism in each case.

For MMP9, the network rule is `proMMP9 AND NOT TIMP1`: proMMP9 promotes activation and TIMP1 suppresses it. With increasing TGF$\beta$, proMMP9 saturates earlier than TIMP1 (top row of Fig. S3), so there exists an intermediate TGF$\beta$ window in which proMMP9 is already high but TIMP1 has not yet reached its plateau. During that window, activation outpaces suppression, producing the MMP9 peak. Once TIMP1 catches up, it clamps MMP9 back down. At high tension, both proMMP9 and TIMP1 are already saturated by mechanical input alone, so MMP9 becomes independent of TGF$\beta$

and remains flat across all doses.

For proMMP1, the analogous decomposition (bottom row of Fig. S3) involves two upstream boosters (NF$\kappa$B and AP1) and one upstream brake (SMAD3). With increasing TGF$\beta$ dose, NF$\kappa$B, AP1, and SMAD3 all rise. Below the TGF$\beta$ level of $\sim 0.42$ (vertical dotted line), both boosters continue to increase and proMMP1 activation rises. Above $\sim 0.42$, the boosters saturate while SMAD3 keeps rising, and the increasing brake contribution causes proMMP1 to decline. As with MMP9, the dose dependence is lost at high tension because mechanical input alone saturates the upstream regulators.

### 9 Cross-direction validation using the dorsal-ventral loading direction

The coupling framework presented in Section 6 was calibrated against the growth stretch measured along the caudal-rostral loading direction. To assess whether the framework generalizes to a second direction, we repeated the calibration against the porcine tissue-expansion dataset using the growth stretch measured along the dorsal-ventral direction. The same equations (Eqs. 4–7 of the main text), the same five coupling parameters ($k_a$, $k_b$, $k_c$, $k_d$, and $\tau_g$), and the same Blackbox optimization procedure (Section 6) were used, with the only change being the experimental target trajectories.

The optimizer was run for 1,315 evaluations, reaching a best loss of 4.97 on the dorsal-ventral target. The calibrated values are

$$k_a = -65.05, \qquad k_b = -51.59, \qquad k_c = -52.10, \qquad k_d = 1.65, \qquad \tau_g = 83.14\,\text{h},$$

yielding the same qualitative structure as the G1 calibration: negative $k_a$ and $k_b$, positive $k_c$ when paired with a small positive $k_d$, and $\tau_g$ on the order of 80–95 h. The model reproduces the temporal increase in growth stretch along the transverse direction at 24, 72, and 168 h (Fig. S4A–C) and yields a directional growth field consistent with the experimental measurements (Fig. S4D–E). The optimization runner (`coupling_G2_optimizer_BB_angles_perTheta.py`), the loss trajectory (`optimization_log_G2_BB_all.csv`), and the figure-generation notebook (`model_predictions_perTheta_wG2.ipynb`) are included in the code repository.

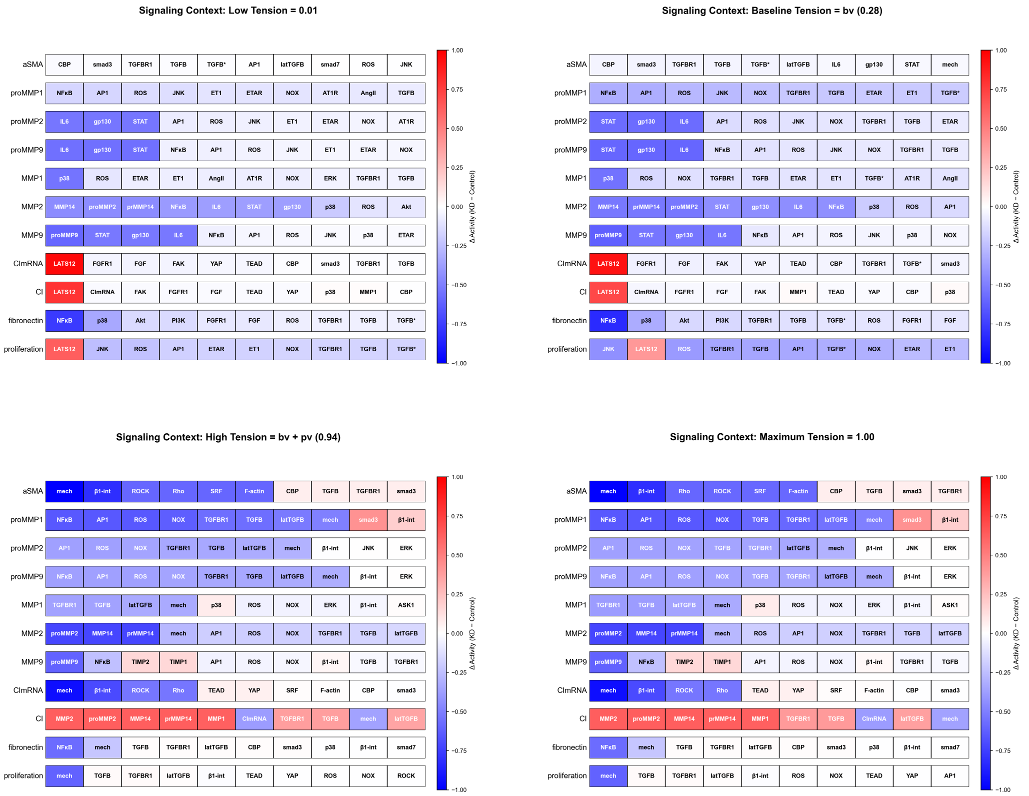


Figure S2: Top-10 most influential knockdowns per phenotype at four tension levels: low (A), baseline (B), high (C), and maximum (D). Each row corresponds to one phenotypic node; columns show the ten knockdowns producing the largest absolute change in steady-state activity, ordered left-to-right by decreasing magnitude. Cell color encodes the change in activity relative to the unperturbed baseline at the corresponding tension level; red (positive) indicates the readout is upregulated upon knockout of the column node (brake), blue (negative) indicates downregulation upon knockout (booster). Self-knockdowns are excluded. Change in activity ranges from −1 to +1.

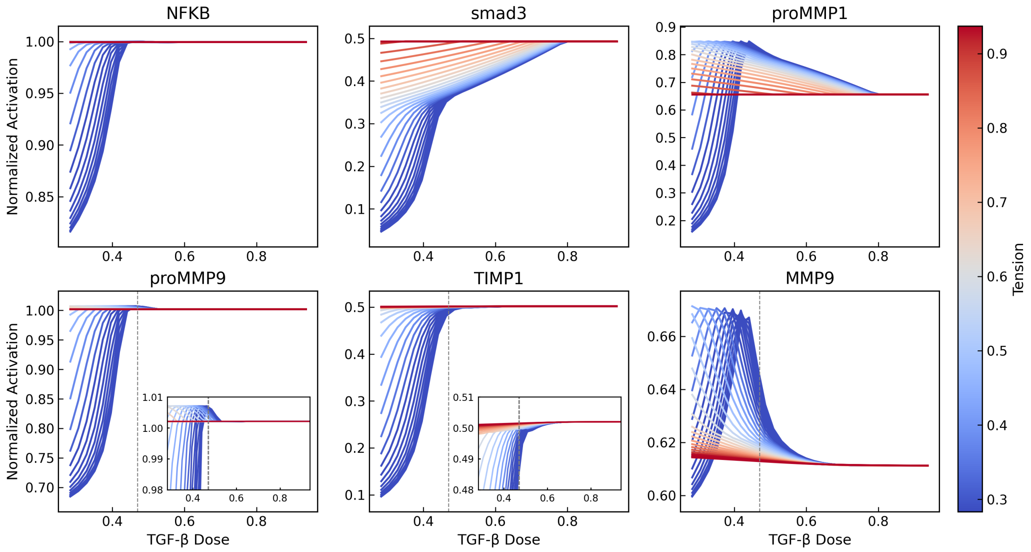


Figure S3: Mechanistic decomposition of non-monotonic dose responses. Top row: proMMP9 activates earlier than TIMP1 with increasing TGF$\beta$; the gap where proMMP9 is high but TIMP1 is still low produces the MMP9 peak in the rightmost panel. Once TIMP1 catches up, MMP9 declines. At high tension, both upstream nodes are saturated independently of TGF$\beta$ and the MMP9 response becomes dose-independent. Bottom row: proMMP1 activation reflects the combined action of two upstream boosters (NF$\kappa$B and AP1) and one upstream brake (SMAD3). Both boosters and SMAD3 rise with TGF$\beta$ until the dose reaches $\sim 0.42$ (vertical dotted line); above that, the boosters plateau while SMAD3 continues to rise, producing the descending proMMP1 limb.

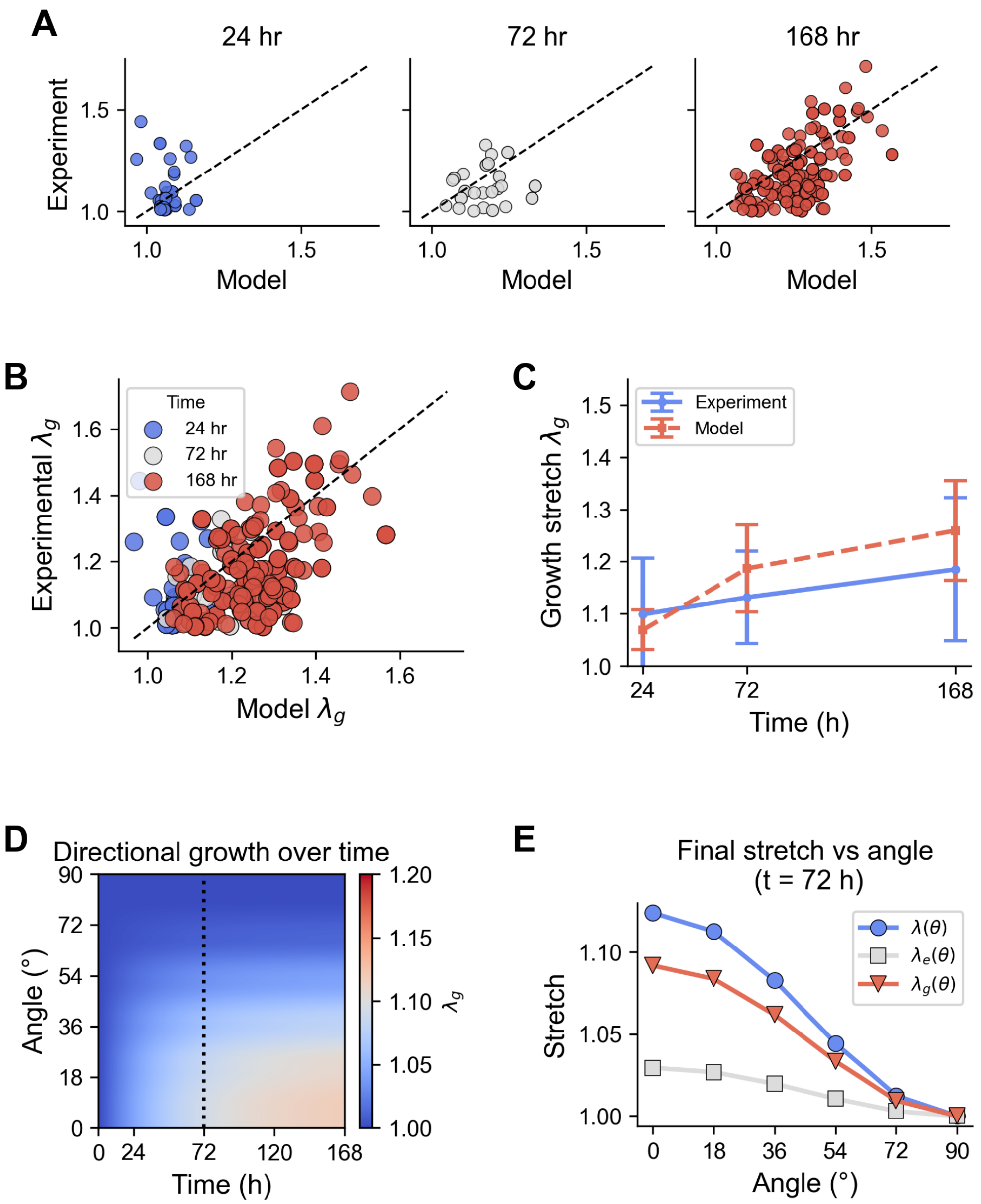


Figure S4: Validation of the coupling framework along the dorsal-ventral loading direction. (A) Comparison between model predictions and experimental measurements of growth stretch $\lambda_g$ at 24, 72, and 168 h. (B) Combined scatter comparison across all samples and time points. (C) Mean and standard deviation of experimental and predicted growth stretch over time. (D) Predicted evolution of directional growth $\lambda_g(\theta, t)$. (E) Angular decomposition of total stretch $\lambda(\theta)$, elastic stretch $\lambda_e(\theta)$, and growth stretch $\lambda_g(\theta)$ at the $t = 72$ h time point. All panels use the same conventions and color coding as Fig. 7 of the main text.
